# Understanding natural selection in Holocene Europe using multi-locus genotype identity scans

**DOI:** 10.1101/2023.04.24.538113

**Authors:** Devansh Pandey, Mariana Harris, Nandita R. Garud, Vagheesh M. Narasimhan

## Abstract

Ancient DNA (aDNA) has been a revolutionary technology in understanding human history but has not been used extensively to study natural selection as large sample sizes to study allele frequency changes over time have thus far not been available. Here, we examined a time transect of 708 published samples over the past 7,000 years of European history using multi-locus genotype-based selection scans. As aDNA data is affected by high missingness, ascertainment bias, DNA damage, random allele calling, and is unphased, we first validated our selection scan, *G12*_*ancient*,_ on simulated data resembling aDNA under a demographic model that captures broad features of the allele frequency spectrum of European genomes as well as positive controls that have been previously identified and functionally validated in modern European datasets on data from ancient individuals from time periods very close to the present time. We then applied our statistic to the aDNA time transect to detect and resolve the timing of natural selection occurring genome wide and found several candidates of selection across the different time periods that had not been picked up by selection scans using single SNP allele frequency approaches. In addition, enrichment analysis discovered multiple categories of complex traits that might be under adaptation across these periods. Our results demonstrate the utility of applying different types of selection scans to aDNA to uncover putative selection signals at loci in the ancient past that might have been masked in modern samples.

## Introduction

With the emergence of large sample size sequencing data, numerous population genetic studies have attempted to identify targets of natural selection in the human genome^1^. However, the majority of studies carried out on modern human populations have largely been restricted to detecting selection events that have happened in the most recent periods of human history because selective sweeps decay due to processes including recombination and mutation ^1^ and can be obscured by demographic events such as admixture^1,2^. By directly tracking genomic changes over time using aDNA, it may be possible to observe sweeps that otherwise are undetectable from modern data. However, until recently, the large sample sizes required for such analyses were unavailable and, as a result, many aDNA based studies to examine natural selection were largely confined to specific alleles^3–7^.

Recently, increased sample sizes have enabled genome-wide selection scans on aDNA ^4,5,8–13^. However, most current approaches have focused on single site statistics that leverage temporal data to detect allele frequency changes over time. An alternative strategy is to use haplotype-based approaches, which are sensitive to footprints of selection left behind by hitchhiking of linked alleles with adaptive alleles. Haplotype scans do not require temporal samples and instead only require samples from a single population group from one specific time to infer recent selective events^14–18^ and might provide complementary information to approaches that detect allele frequency changes over time. However, most haplotype-based methods require phased genomes that are particularly challenging to obtain from ancient samples for several reasons. First, aDNA read lengths are incredibly short (30-50bp) and read-based phasing has reduced efficiency at these lengths^19^. Second, reference panels constructed from modern haplotypes may introduce bias in calling alleles from aDNA due to divergence that has arisen between ancient and modern genomes. By using trio or family data, where the phasing and imputation can be assessed directly and precisely by examining transmission of alleles, biological information can be used to obtain a ground truth dataset. However, due to the nature of sampling in aDNA studies, there are relatively few trios or families that have been sequenced of sufficient quality that may help with assessing the quality of phasing and imputation methods.

Recently, statistics that leverage multi-locus genotypes, which represent strings of unphased genotypes from diploid individuals, were proposed to circumvent the need for phased haplotypes^20–22^. However, a major challenge in applying these statistics to aDNA is its low coverage (largely between 0.5-2x coverage), which results in, on average, only one of the two diploid alleles being called. Moreover, the reference allele in modern genomes may bias which of the two diploid alleles is mapped. In this study, we modified a multi-locus genotype-based scan^22^ for adaptation to be suitable for low-coverage aDNA data using a pseudo-haploidization scheme, in which one allele per site is randomly selected to represent the genotype of the individual at that position. We evaluated the performance of this method, which we call *G12*_*ancient*_, on aDNA using simulations and well characterized functionally validated variants. We then applied it to different epochs from an aDNA time transect to examine the timing of selection of well-characterized candidate sweeps. Finally, we examined novel targets of selection to see if our new method could complement other studies of natural selection carried out using allele frequency-based methods^13^.

We carried out this analysis on a dataset of ancient individuals from Holocene Europe representing a period of significant cultural change, beginning with the transition from hunting and gathering to farming, which resulted in people living in much closer proximity to animals, as well as major dietary changes. This was also a period that covered the transition to state-level societies, which led to large population densities and division of labor^23^. Notably, several papers document the first evidence of bacterial and viral pathogens in the aDNA record during the Holocene, and it is of interest to understand if and how humans adapted to these new cultural changes, environments, and diseases that affected us in our evolutionary past^10,24,25^. Given the large sample sizes spanning this time transect that provide a nearly gapless record of human populations in Europe, we attempted to estimate the timing of selection and generate hypotheses about its correspondence with major demographic and cultural changes.

## Results

### A time transect through Holocene Europe

In our analysis we examined a collection of 708 recently published samples from Europe ranging from 6572 BP to 353 BP (**Supplementary Data**)^26–38^. To minimize reference bias or batch effects associated with data processing issues across the set of samples, we chose to include only samples for which hybridization capture was performed on 1.2 million positions^39^ and that had at least 15,000 SNPs for which we could perform high-quality population genetic analysis. We only included samples that did not appear to have significant contamination on the mtDNA or the X chromosome (in males) and were unrelated (up to the third degree). We also chose to only include samples that were uniformly treated with the same Uracil-DNA Glycosylase (UDG) process during library preparation and trimmed the last two bases from each read to reduce the impact of aDNA damage on our computed statistics (**Methods:** *aDNA data curation*).

To homogenize the sample size of our analysis across time periods, we used 177 individuals for each epoch which we determined based on *f*_*4*_-statistics, time period (based on direct radiocarbon dates or precisely dated archaeological contexts), and geographic location (**Fig. 1**). Samples from each of these assigned population groups were genetically homogenous and had little to no ancestry from additional sources known to enter Europe and contributed in small proportions to a minority of European populations, including the Scythians and Sarmatians, the Uralic-related migrations into Hungary and Fennoscandia, and Iranian farmer related ancestry along the Mediterranean in Southern Europe. The groups of individuals were:

**Fig. 1:**
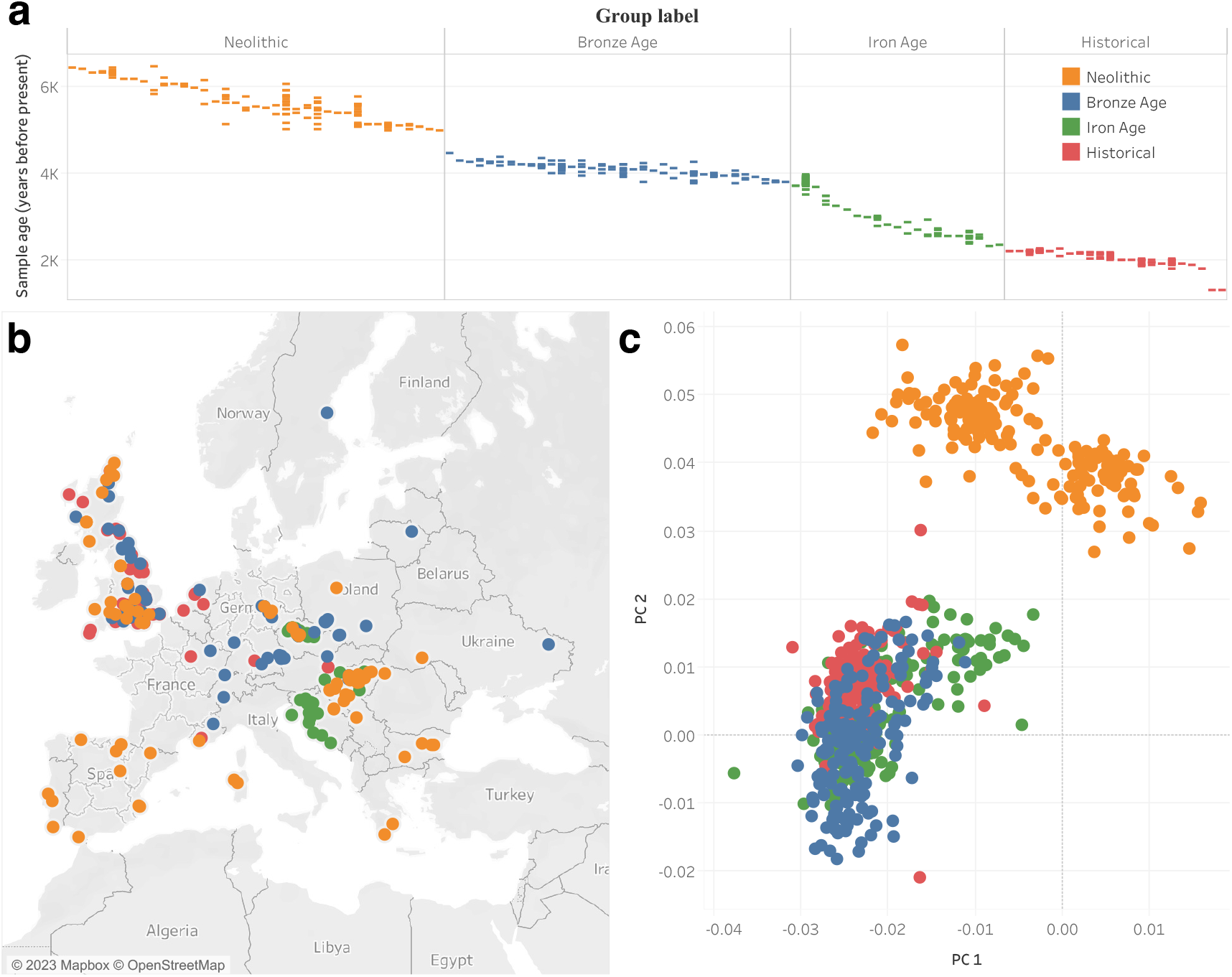
aDNA samples included in this study. **a** Distribution of archeological or radiocarbon dates for sites (vertical columns) from each time period over the past 7,000 years. Each colored bar represents single samples from a site that has been dated to a particular time. Multiple samples from the same site are annotated along the same column. **b** Locations of ancient individuals that passed our analysis thresholds, forming a sample size of 708 individuals. **c** PCA analysis of ancient individuals projected onto a basis of modern samples.

**N** First farmers of Europe from the Middle to Late Neolithic (abbreviated as the first letter of <u>N</u>eolithic). These individuals were from across Europe, are dated to between 6572 and 5091 BP and are mixtures of European Hunter-Gatherer and Anatolian farmer ancestry.

**BA** Bronze Age Europeans (abbreviated as the first letters of <u>B</u>ronze <u>A</u>ge). These individuals are from the Bell Beaker cultures of Western and Central Europe dated between 5940 to 3780 BP.

**IA** Iron Age Europeans (abbreviated as the first letters of <u>I</u>ron <u>A</u>ge). We used samples from Iron Age Britain and other countries in Western Europe dated between 3465 to 2130 BP.

**H** Finally, to represent <u>H</u>istorical samples from Europe, we included samples from the Roman and late antique periods, primarily from Britain, dated from 1973 to 353 BP.

### A modified multi-locus genotype statistic for detecting selection on aDNA

For application to unphased data, several multi-locus genotype methods have been recently developed that are similar to extended haplotype-based statistics^21,22^. Evidence based on simulation studies have suggested that these approaches using unphased information might be as powerful as approaches that use phased information^21,22^. However, the low coverage (mean: 1.5×) of aDNA data means that we are unable to call heterozygotes effectively and are therefore unable to use these statistics directly. Due to this low coverage, aDNA samples are processed as ‘pseudo-haploid’ data where one read mapping to a position is chosen at random and the allele of that read is used as the genotype (pseudo-haploidization) (**Supplementary Fig. 1** and **Methods:** *Generation of modern human data mimicking ancient data*).

To examine selection on this type of data, we adapted an approach that has been previously shown to be useful in application to unphased population genomic data, *G12. G12* is capable of detecting selective sweep signatures associated with hard sweeps, expected when adaptation is gradual, and soft sweeps, expected when adaptation is rapid^40,41^. We modified *G12* to work on pseudo-haploidized aDNA data. This modified statistic which we call *G12*_*ancient*_ is computed in windows comprising a fixed number of SNPs and is defined as:

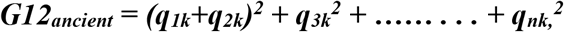

Where *q*_*1k*_, *q*_*2k*_, *q*_*3k*_, ……., *q*_*nk*_, denote the frequencies of the unique n, pseudo haploidized multisite genotypes, ranked from most common to most rare. The intuition behind this statistic is that haplotypes that have risen to high frequency are likely to have a large number of individuals with homozygous genotypes (thereby biologically phased as the two haplotypes are identical) and that these homozygous haplotypes provide a similar signal to those from phased data.

To validate our modified statistic and its applicability to aDNA data, we took several approaches. As a first line of analysis, we examined the correlation between *G12* computed on diploid low-coverage data from the 1,000 genomes project^42^ with that of *G12*_*ancient*_ computed on the same samples using a pseudo-haploid genotyping scheme along with introducing missingness and ancient DNA damage at typical rates in our dataset (**Methods:** *Running selection scans on ancient datasets* and **Supplementary Fig. 3**). The correlation between *G12* and *G12*_*ancient*_ across all windows in the genome was 0.95 suggesting that our modified version of the selection statistic *G12*_*ancient*_ is almost equivalent to running *G12*, a selection statistic that has already been examined previously and applied to various other datasets. (Methods: *Running selection scans on ancient datasets* and **Supplementary Fig. 3**). Second, we tested the ability of *G12*_*ancient*_ in simulated data to demonstrate its performance on a range of theoretical settings. Third, we demonstrate the ability of *G12*_*ancient*_ to identify well-characterized and functionally validated variants that have previously been found to be under selection by multiple modern and ancient genomic studies^8,43^ (**Supplementary Table 2**).

### Evaluating G12_ancient_ on simulated data

To evaluate the performance of *G12*_*ancien*t_ in simulated aDNA data, we used the forward in time simulator SLiM 3^44^ to generate genotypes incorporating missingness, ascertainment bias, random allele calling, and genotyping error (**Methods:** *Generation of simulated data*) that are typical of the aDNA data used in our study. We simulated hard and soft sweeps in a population under the Tennessen et al. model^45^, a demographic model that captures broad features of the allele frequency spectrum of modern European genomes. We varied the time of the onset of selection, the selection coefficient (*s*), and the time period of the sample. We obtained three samples of 177 individuals, matching the sample size of our dataset, spanning the past ∼7,000 years (250, 100 and 40 generations before present).

We first showed that our pseudo-haploidization approach does not reduce the ability of *G12*_*ancient*_ to detect selection, and that the distribution of *G12*_*ancient*_ values of pseudo-haploidized simulated data is comparable to that of running the haplotype-based statistic *H12* on phased data (**Supplementary Fig. 4**). When incorporating missingness and data sparsity at levels typically observed in aDNA to our simulated datasets (**Methods:** *Running selection scans on simulated data*) the *G12*_*ancient*_ signal is attenuated but can still be differentiated from neutrality. (**Fig. 2** and **Supplementary Fig. 5**). Additionally, we observe that G*12*_*ancient*_ increases with stronger selection (**Fig. 2** with the exception of **Fig. 2**. bottom row). In all selection scenarios analyzed, with the exception of young sweeps with weak selection, selection can be easily distinguished from neutrality (s = 0).

**Fig. 2:**
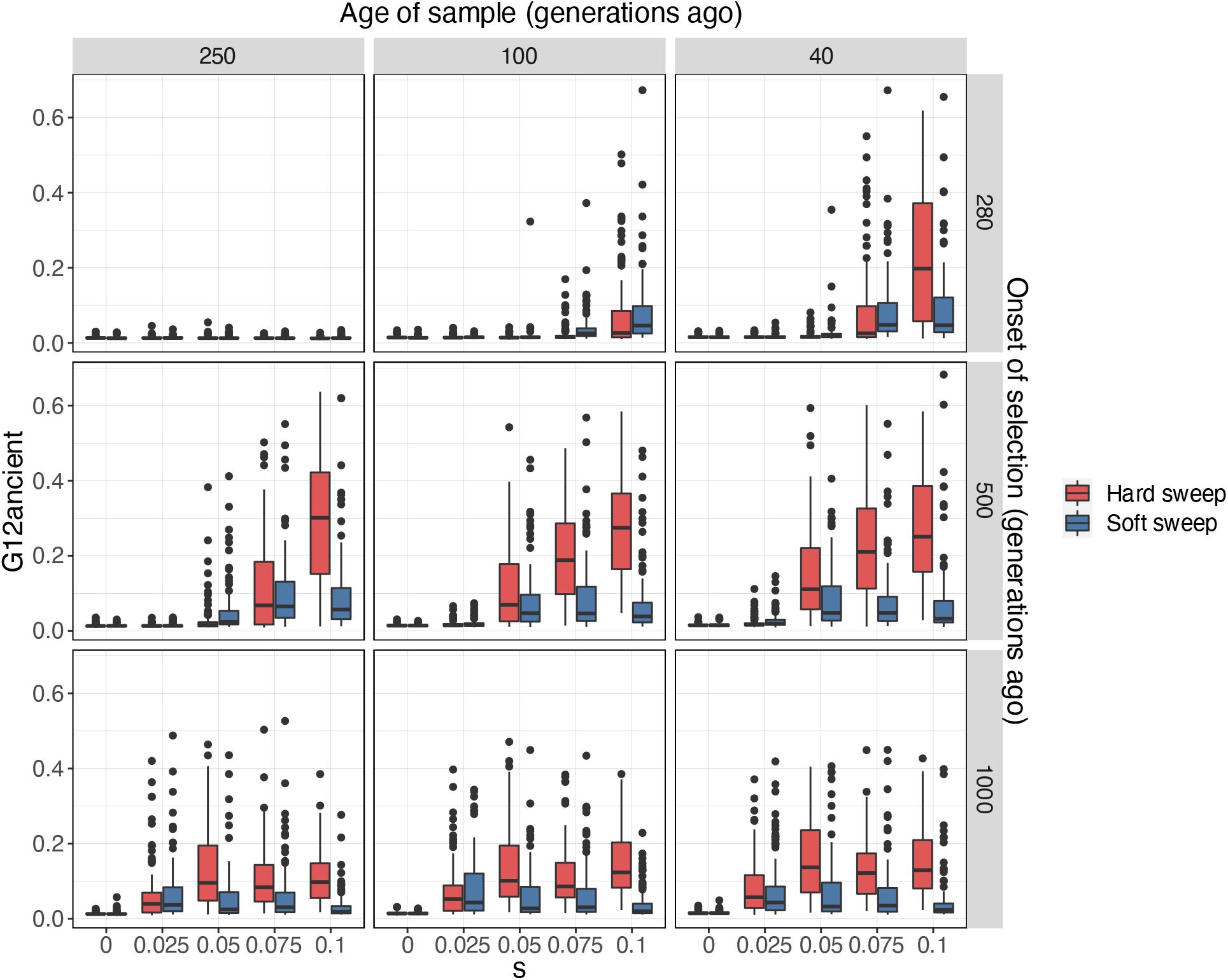
G12_ancient_ values for hard (red) and soft sweeps (blue) in simulated aDNA data. *G12*_*ancient*_ values were obtained for varying selection coefficients (*s*) and onset of selection (rows). We sampled the population at 3 different time points (columns). For the soft sweep simulations *K*=25 independent adaptive mutations were introduced to the population at the time of the onset of selection. We ran a total of 100 simulations for each combination of parameters with mutation rate μ = 1.25×10^−8^ /bp, chromosome length L=5×10^5^ and recombination r = 1×10^−8^ events/bp. Selection *s* = 0 represents neutrality.

In addition, we assessed the ability of *G12*_*ancient*_ to detect sweeps of varying degrees of softness. To do so, we introduced *K* beneficial mutations at the time of the onset of selection for *K=5, 10, 25* and *50* (**Supplementary Fig. 6**). For *K* = 5 the majority of the resulting sweeps were hard, whereas for higher values of *K* the probability of a sweep being soft increased (**Supplementary Fig. 7**). Again, *G12*_*ancient*_ was able to distinguish selection from neutrality for varying degrees of softness except for sweeps that were very young (**Supplementary Fig. 6** first row, sample from 250 generations ago). Additionally, we observed that as sweeps became softer, the *G12*_*ancient*_ signal decreased, making it harder to detect sweeps that are old and very soft (**Supplementary Fig. 6** bottom row, *K* = 50).

### Application of G12_ancient_ to functionally validated variants from real data

To test the ability of *G12*_*ancient*_ to detect selection signals on real data, we modified modern genomic data from European individuals from the 1000 genomes project^42^, by introducing missingness, ascertainment bias, sample size and random allele calling to mimic aDNA (**Methods:** *Generation of modern human data mimicking ancient data*). We then examined the ability of *G12*_*ancient*_ to detect classic selective signals in the genes *LCT, TLR1* and *SLC24A5* which have been identified by multiple previously conducted selection scans and are regions that are highly differentiated between Europeans and Asians (**Supplementary Table 2**). The causal alleles at these loci have been fine mapped in association studies and have also been functionally validated in cellular assays. These alleles are commonly used as positive controls in studies carrying out tests for natural selection in humans^43^. The *LCT* locus is responsible for conferring lactase persistence into adulthood; *TLR1* is a gene involved in immune cell response and *SLC24A5* is the dominant locus contributing to light skin pigmentation in Europeans^3,46^.

Using our aDNA mimicking process on the modern data and then applying *G12*_*ancient*_, we were able to identify the *LCT, SLC24A5* and *TLR1* loci in the top 3 peaks observed chromosome-wide in the European (CEU) population but not African (YRI) and South Asian (STU) populations (**Fig. 3a**). We also examined the effect of utilizing different parameters for window-sizes and jumps (distance between analysis windows) and obtained an optimal choice of these parameters on real data (**Methods:** *G12ancient parameter choices and peak calling* and **Supplementary Fig. 8**).

**Fig. 3:**
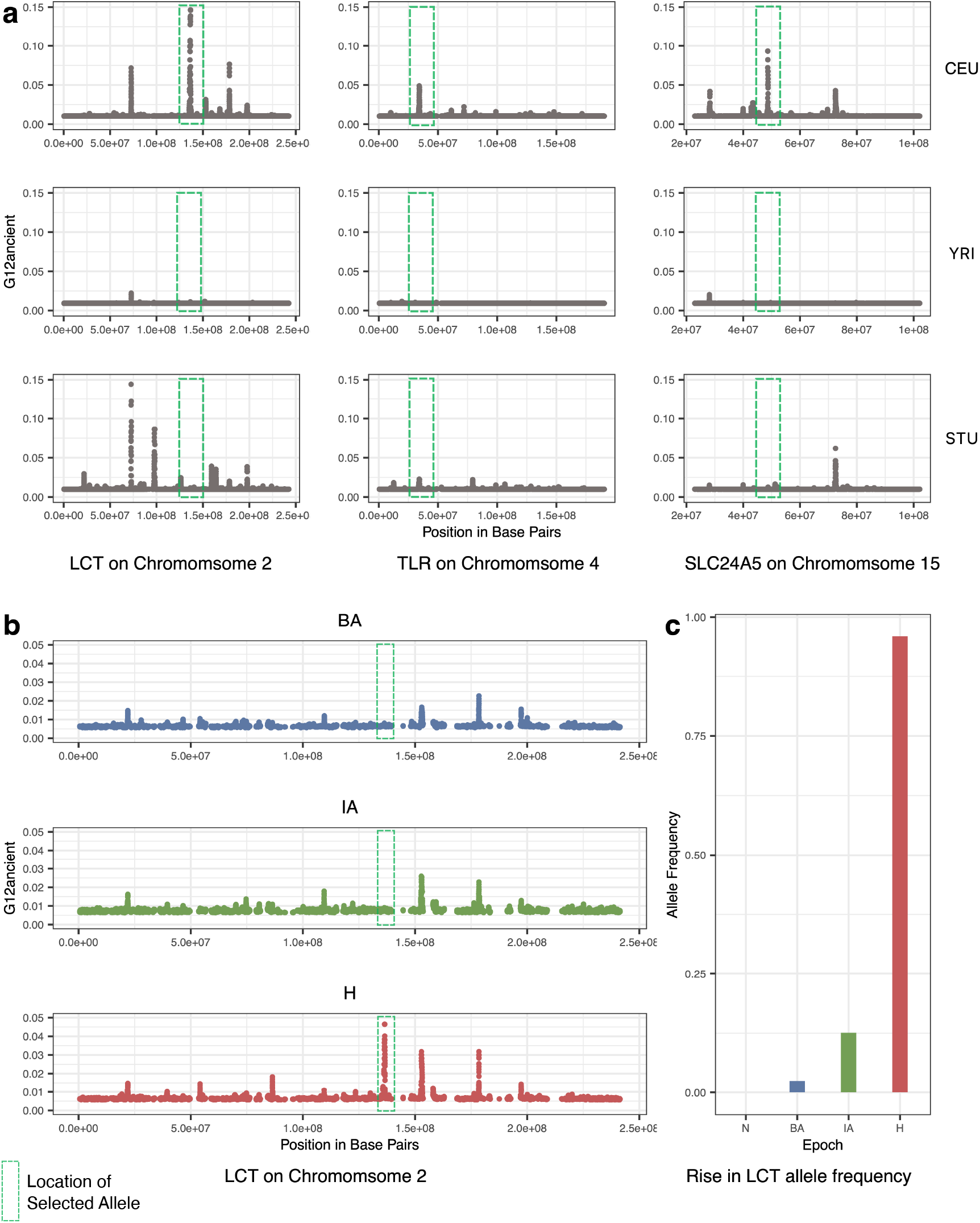
Recovery of variants well characterized to be under selection in modern Europeans (positive controls). **a** *G12*_*ancient*_ values for modern population data from the 1000 genomes project^42^, which was modified to mimic aDNA. *G12*_*ancient*_ can detect several variants that have been previously found to be under selection in modern Europeans. However, *G12*_*ancient*_ is completely absent or highly attenuated in populations of other ancestries (YRI and STU). **b** Using aDNA data, it is observed that the signal for the *LCT* allele is absent in BA and IA populations but is a top peak genome-wide in the H population. **c** The allele frequency reaches near fixation in the H population but is absent in N period as it was only introduced into Europe by the arrival of pastoralists from the Pontic-Caspian Steppe^13^. In panel **b** we show that we observe high *G12*_*ancient*_ only in the historical period but not in previous time periods as a demonstration of our ability to localize the timing of selection to various epochs.

Next, to establish that our process could identify the timing of signals of natural selection from aDNA, we examined the *LCT* locus at different time periods of European history. This locus is particularly relevant for this analysis as the causal allele was absent in Europe prior to the arrival of Steppe Pastoralists in the Bronze Age and, therefore, could not have been under selection prior to that point^3,8,12,13,43,47–51^. By applying *G12*_*ancient*_ across different periods in our time transect, we show that we were able to identify selection at the *LCT* locus, in the historical period (this window is the top peak genome-wide), but we do not see signals of selection for these in the Bronze Age and Iron Age populations (**Fig. 3b**). These results therefore are in line with the rapid increase in frequency of the causal variant rs4988235 only in the historical period (**Fig. 3c**), a finding that has also been replicated in other aDNA studies^8,13,43^.

### Time stratified selection in ancient Europe

Having established that our selection scan can identify signals of selection in simulated data and correctly distinguish between positive and negative controls in real data, we applied *G12*_*ancient*_ to the aDNA time transect. We defined a genome-wide threshold for significance as the 5th highest *G12*_*ancient*_ value obtained by simulating neutral data under the Tennessen et al model^45^ (**Methods:** *Running selection scans on simulated data, G12ancient parameter choices and peak calling*) *G12*_*ancient*_ values above this threshold were classified as putative sweeps. As windows adjacent to each other may belong to the same selective sweep, consecutive analysis windows above the *G12*_*ancient*_ neutral threshold were assigned to a single peak. The highest *G12*_*ancient*_ value among all windows of a peak was chosen to represent the whole peak. To remove spurious peaks that might have arisen due to high rates of missing data or low recombination rates, we masked the peaks located in those regions (**Methods:** *Quality control for removing false sweeps*). With this approach, around 3-4 peaks per epoch were obtained that reached significance at the genome-wide level.

We began by re-examining 12 loci previously established to be under selection using aDNA data^8^. Although the selection signals produced by the previous scan and our scan differ in their methodology and, therefore, their ability to detect selective events, we wanted to assess if we might be able to use our approach to localize when in time these signals were selected.

As seen in **Fig. 4a**, the time period in which we observe a signal of selection at the *LCT* locus is limited to the historical period. In the N population, among the top peaks, we found a signal which included the gene *OCA2*/*HERC2*, variations in this gene are associated with eye, skin, and hair pigmentation variation^8,26,52^. This gene is the primary determinant of light eye color in Europeans and in our analysis, we time the selection signal to the Neolithic^13^. However, we observe a signal of selection at the *HLA* and at neighboring *ZKSCAN3* in most of the epochs (**Fig. 4a** and **Fig. 4c**).

**Fig. 4:**
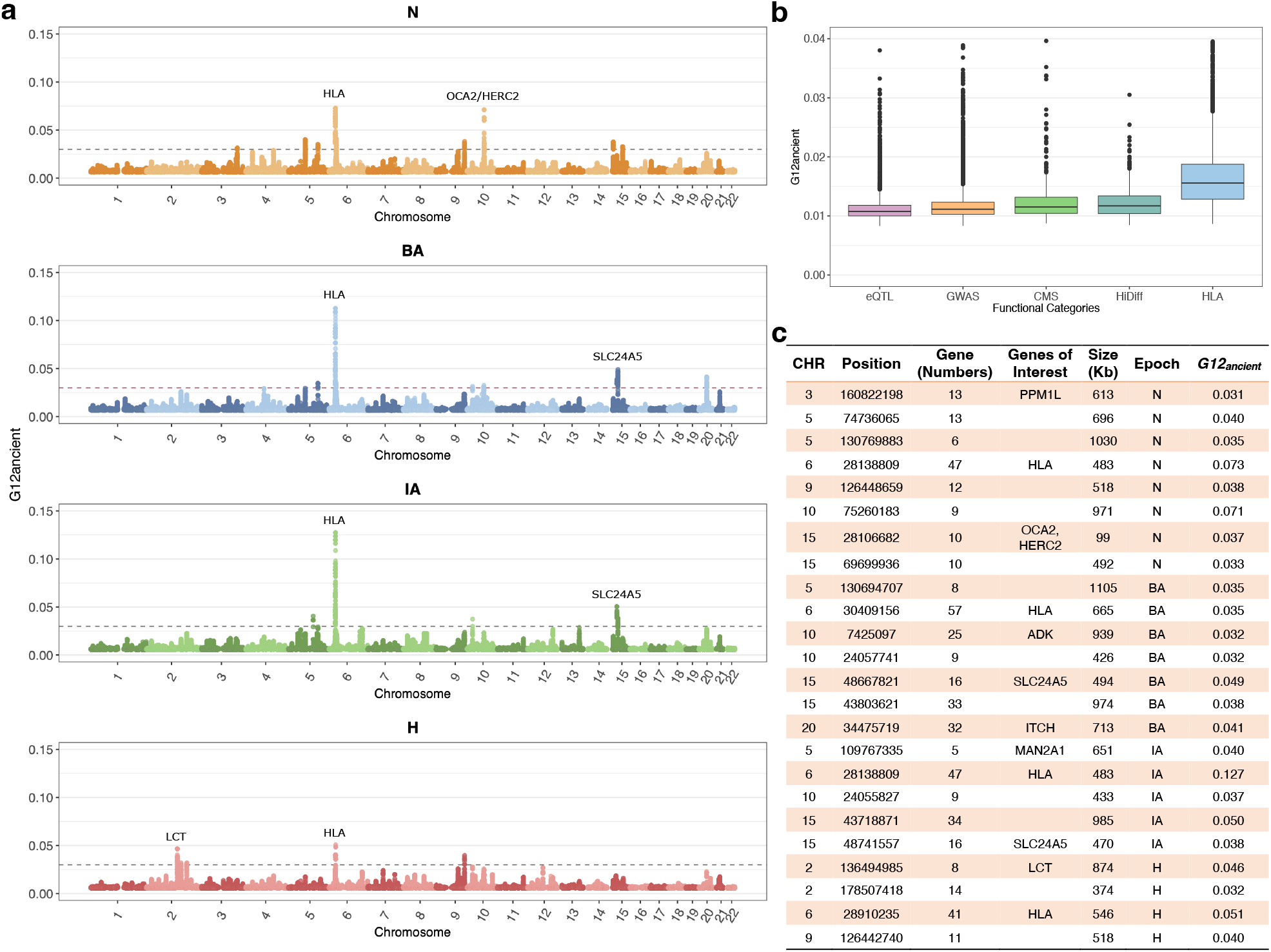
G12_ancient_ applied on aDNA data. **a** Manhattan plot of *G12*_*ancient*_ values genome-wide with the top signals in each epoch annotated. The gray dashed line is the genome-wide significance threshold based on simulations under the Tennessen et al. demographic scenario^45^. **b** Boxplots showing the variation of *G12*_*ancient*_ across various functional categories. **c** Signals from genome wide significant top scoring loci for different epochs. The column Gene (Number) represents the number of genes mapped to the peak. For some genes, we assign a gene of interest based on fine mapping studies that have examined the results of modern selective sweeps examining the same regions.

Outside of these 4 loci, our selection scan also revealed several other candidates which we determined as being above our significance threshold. Several of these were associated with skin and eye pigmentation. In the BA and IA epochs we observed a signal of selection in the gene *SCL24A5*. As mentioned above, this gene is thought to be the major determining locus for light skin pigmentation in Europeans^3,53^. While highly differentiated between Asians and Europeans and appreciated as a major candidate of selection using modern European Genomes, single SNP allele frequency approaches examining aDNA have yet to identify this particular allele as a candidate^8,13^. This shows the value of employing alternate types of selection scans on similar datasets to uncover putative selective sweeps.

We observed a signal at a locus associated with *PPM1L* as on the top peaks in N, which is an obesity related marker in Humans^54^. This signal for selection on obesity and body weight related alleles during the Neolithic or the change in dietary practices from hunting and gathering to farming is also observed in single SNP based approaches^13^.

We also observed several signals in genes that were associated with immunity or auto-immunity. In the BA population, we observed a candidate in locus containing *ADK*, which regulates the intra and extracellular concentrations of adenosine which has widespread effects on cardiovascular and immune systems^55,56^. We see a signal at the *ITCH* gene in the BA, which is associated with immune response, and regulation of autoimmune diseases^57,58^. In the IA we see candidate sweep at the *MAN2A1* locus - genetic variations in this gene have been shown to cause human systemic lupus erythematosus^59^. In **Fig. 4c** we report a list of all regions that appear to be under selection in each epoch along with some genes of interest that lie in those regions.

### Gene Set Enrichment Analysis

In addition to examining individual SNPs, we examined mean *G12*_*ancient*_ values across broad categories of functional SNPs. We looked at loci that were associated with changes in gene expression (eQTLs), identified as associated in Genome Wide Association Studies (GWAS), or were thought to be previously under selection in Europeans (CMS) or highly differentiated between Europeans and Asians (HiDiff) or were part of the HLA region. We found that functional categories of SNPs were seen with significantly higher *G12*_*ancient*_ values compared to SNPs that were not annotated as being functionally relevant, with the HLA region being the most elevated of the functional categories (**Fig. 4b**).

We next asked if we could associate biological functions to these top-scoring loci. We computed a p-value based on deviation from neutrality based on simulations (**Methods:** *Enrichment Analysis*). To determine if categories of genes associated with Genome-Wide Association Studies were significantly associated with selection signals, we carried out enrichment analysis using FUMA^60^, which maps SNPs to genes and performs gene set enrichment analysis for GWAS annotations incorporating LD information as well as gene matching by length and conservation scores (**Methods:** *Enrichment Analysis)*. We found that many categories of GWAS related to anthropometric traits, auto-immune traits as well as disease related traits were under selection across the different time epochs. We report gene sets from the GWAS catalog using FUMA: Gene2Func^60^ and used a conservative significance threshold of – log_10_ *p* >= 5. We report the list of all categories for which we observed enrichment in **Fig. 5**.

**Fig. 5:**
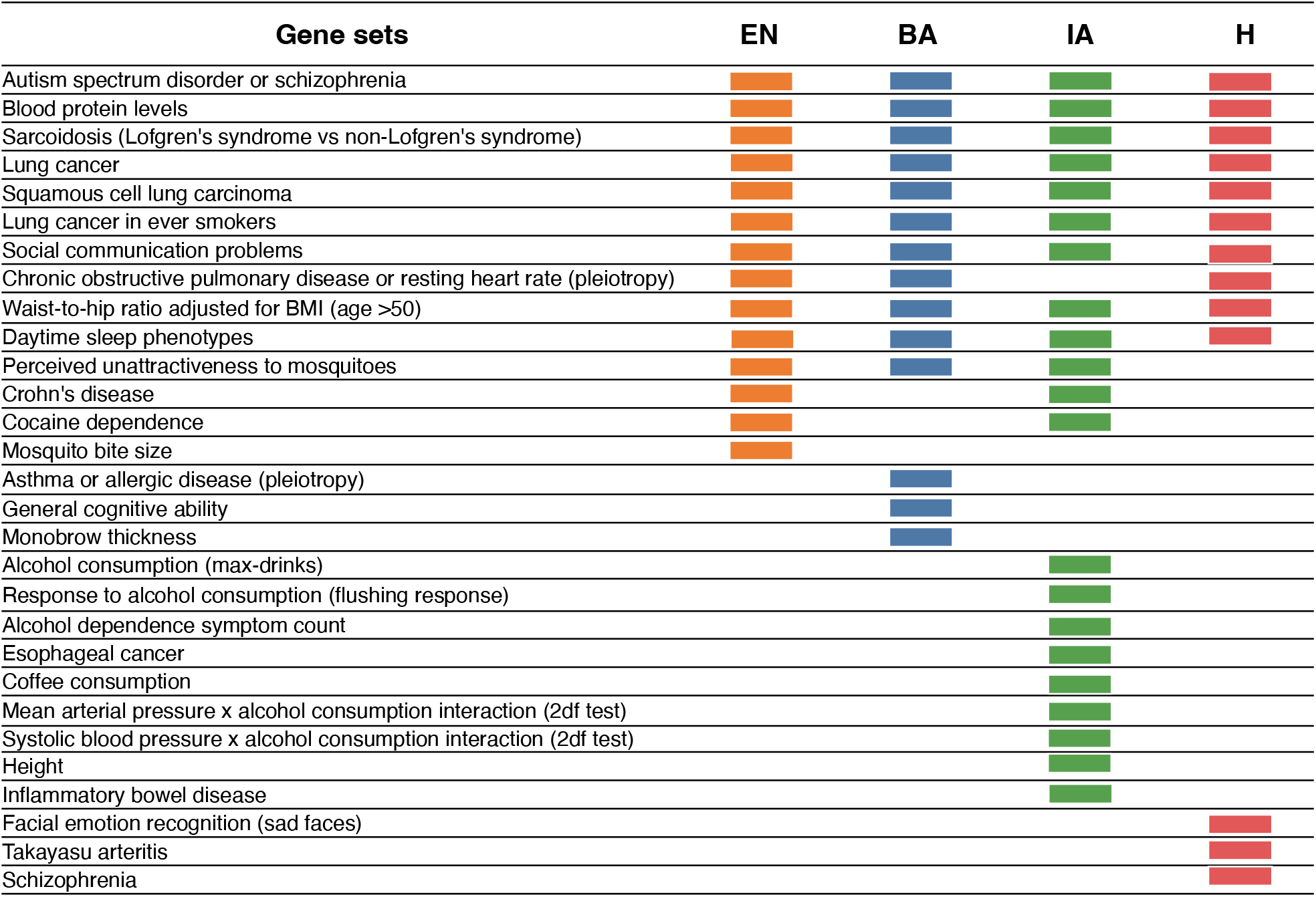
Gene sets enriched across epochs. Colored boxes show significantly enriched gene sets for each epoch. Several gene sets are enriched across the 4 time periods.

## Discussion

In this paper, we introduce a modified version of a previously described selection statistic^16,22^ and applied it to a time transect of aDNA from Europe. To date, while allele frequency-based approaches have been used extensively in the field, approaches using haplotype scans have largely been lacking. A single study^61^ performed a selection scan by phasing low coverage aDNA samples, and running a widely used extended haplotype statistic, XP-EHH. Here, we took an alternate approach aimed to reduce bias and artifacts from the use of modern reference panels for phasing and imputing low coverage ancient DNA, but largely maintaining power when compared to phased approaches in simulations.

Our results, which take advantage of the major increases in sample size in the availability of aDNA data in the past 5-10 years demonstrate the potential of running multi-locus genotype-based scans on aDNA. Our modified statistic, which we verified through simulations and gold standard variants, can potentially be employed in other settings where sequencing coverage is low and there is high missingness requiring pseudo-haploidization. Importantly, since haplotype-based statistics are not as reliant on temporal data to exclude false positives, these statistics are useful for ancient datasets from geographic regions that only have a single timepoint.

Despite its potential, our approach also has several limitations. As the results from the simulation study show, our statistic is powered mostly for strong selective sweeps (*s* > 0.01). Moreover, the timing of onset of selection is limited by our ability to detect selection below this high threshold and therefore lack of selection at a particular time could also be due to a lack of power. Another major limitation of our approach is that our window-based method is unable to localize selection to a specific allele as it is based on detecting deviation in a surrounding region of 200 SNPs. On data from a capture array like we have, this distance can span large distances and decrease our target resolution. Here we used the closest gene to the peak SNP in a series of windows to connect genes to candidates under selection.

An important future direction for this type of research is to carefully examine the accuracy of imputation and phasing on low-coverage ancient data using biological confirmation such as from trios, which could become more available as coverage increases for many samples due to lower sequencing cost and better technology. Additional directions could also be to extend these scans to other time periods or more importantly to other geographic regions in the world where aDNA data is becoming rapidly available.

## Methods

### aDNA data curation

We obtained aDNA data from Allen Ancient DNA Resource^62^ (https://reich.hms.harvard.edu/allen-ancient-dna-resource-aadr-downloadable-genotypes-present-day-and-683, version 51), and selected the samples that were enriched for 1240k nuclear targets with an in-solution hybridization capture reagent. We did not include individuals if they had less than a 3% cytosine-to-thymine substitution rate in the first nucleotide for a UDG-treated library as these were indications of contamination. We also removed individuals who had indications of contamination based on polymorphism in mitochondrial DNA or the X chromosome in males, based on estimates from contamix^63^ and ANGSD^64^. For population genetic analysis to represent each individual at each SNP position, we randomly selected a single sequence (if at least one was available).

Finally, we assembled genome-wide data of various human populations from Holocene Europe dated between ∼7000 BP and 500 BP. To maintain homogeneity across time periods, we sampled 177 individuals from each archeological period - the Neolithic, the Bronze Age, the Iron Age and Historical period. For populations with more than 177 individuals, we only chose samples from these periods with the highest coverage and prioritized samples from the same site whenever possible. A list of all samples analyzed is in **Supplementary Data**.

### Principal components analysis

We carried out PCA using the smartpca package of EIGENSOFT 7.2.1106^65,66^. We used default parameters and added two options (lsqproject: YES, and numoutlieriter:0) to project the ancient individuals onto the PCA space. We used 991 present-day West Eurasians as a basis for projection of the ancient^31,67^ We restricted these analyses to the dataset obtained by merging our aDNA data with the modern DNA data on the Human Origins array and restricted it to 597,573 SNPs. We treated positions where we did not have sequence data as missing genotypes.

### Generation of modern human data mimicking ancient data

To examine whether the *G12*_*ancient*_ based selection scans would be applicable to aDNA data; we developed a process of converting the modern human genomic data from the 1000 Genomes project^42^ to mimic the statistical and physical properties of aDNA data and ran the scans on modified modern data. We utilized a pseudo-haploidization scheme in which we randomly selected (probability of selection is 0.5) one of the two alleles from the heterozygous genotype as described in **Supplementary Fig. 1**.

To simulate ascertainment, we restricted the 1,000 genomes samples to just the 1.2 million positions that were on the aDNA capture array. Finally, we incorporated missingness on a per site basis in modern data using the mean (0.55) and standard error (0.23) we observed in our sample of 708 individuals and randomly set the genotypes of a certain proportion of individuals in the modern data to missing (**Supplementary Table 1** and **Supplementary Fig. 2**)

### Running selection scans on modern data mimicking aDNA processing

We ran *G12* on 91 GBR individuals from the 1000 Genomes^42^ with phased genotypes called using the standard process and *G12*_*ancient*_ with the same individuals processed using our ancient DNA mimicking approach. We then compared the *G12* values and *G12*_*ancient*_ values at each SNP and calculated the Pearson correlation coefficient between *G12* and *G12*_*ancient*_ values and found out they are strongly positively correlated with each other with a correlation coefficient of 0.95 (**Supplementary Fig. 3**) suggesting that our new statistic behaved similarly to the original *G12* statistic on our data.

### Generation of simulated data

We used SLiM 3.7^44^ to simulate hard and soft sweeps under the Tennessen et al. demographic model^45^ with mutation rate μ= 1.25×10^−8^ /bp, chromosome length L=5×10^5^ and recombination r = 5×10^−9^ events/bp. In the hard sweep simulations, a single beneficial mutation was introduced to the population. The simulations were conditioned on the presence of the adaptive mutation, that is, we restarted the simulation if the adaptive mutation was lost. To model soft sweeps, we added K=5, 10, 25 and 50 distinct copies of a beneficial mutation. We varied the time at which these mutations were introduced, t= 280, 500 and 1000 generations ago, along with their selection coefficient (*s)* and sampled the population at three different time points: 250, 100 and 40 generations before present.

Based on the missingness observed in our ancient DNA data, we added missing data to our simulated datasets following a beta distribution with mean 0.55 per SNP and standard deviation of 0.23 (**Supplementary Table 1**). Moreover, we followed the pseudo-haploidization scheme used in processing the data (**Supplementary Fig. 2** and **Supplementary Fig. 4**). Finally, in order to incorporate the sparsity of aDNA data, we randomly selected 201 SNPs from our pseudo-haplotype data. That is, we obtained a 201 SNP window for our sample of 177 individuals.

### Running selection scans on simulated data

We computed *G12*_*ancient*_ in simulated data using 201 SNP windows in a total of 100 simulations for each combination of parameters tested. We first obtained *G12*_*ancient*_ for hard sweeps and neutrality (*s* = 0) with and without applying our pseudo-haploidization scheme and with no missing data (**Supplementary Fig. 4**). We varied the strength of selection and the time of the onset of selection (age of mutation, in generations).

Next, we obtained the distribution of *G12*_*ancient*_ values in data sets containing missing data. We compared the *G12*_*ancient*_ signal obtained from hard sweep and neutral simulations with and without missing data, obtaining a reduction of signal when missingness was included (**Supplementary Fig. 4**).

We tested the ability of *G12*_*ancient*_ to detect sweeps across various degrees of softness in sparse genomic data with high missingness. We introduced *K* beneficial mutations at the time of the onset of selection for *K=5,10,25* and *50* (**Supplementary Fig. 6**). To determine whether these simulations were more likely to result in hard or soft sweeps, we computed the number of distinct mutational origins at the selected site in each simulation (**Supplementary Fig. 7**). When *K=5* most simulations have a single origin, giving rise to hard sweeps. As *K* increases, so does the number of origins in the simulations, increasing the probability of soft sweeps as well as the softness of the sweep.

### Running selection scans on ancient datasets

After examining the application of *G12*_*ancient*_ on simulated data, we examined our ability to identify 3 major signals of adaptation previously observed in modern Europeans^43^. We list them here along with their known functional impact.

### G12_ancient_ parameter choices and peak calling

To calibrate our *G12*_*ancient*_ statistic we iterated over several parameter choices to improve performance. The most significant parameters are window and jump. Window refers to the analysis window size in terms of SNPs, and jump is the distance between centers of analysis windows (<u>readme.pdf</u>). To find the best combination of window and jump, we ran a grid search and varied the window size from 50-400 SNPs with a step size of 25 SNPs and jump from 1-20 with a step size of 5 SNPs. We tried to optimize our process on the three signals of well characterized adaptation in humans from the previous section on the H population which is closest in time to modern samples. Larger window sizes resulted in decrease of *G12*_*ancient*_ values, and larger step sizes resulted in decrease of SNP density, as larger windows diminish the power of the statistic by averaging over regions that are come from different linkage blocks. As jump increases, fewer and fewer SNPs are used in the computation, as illustrated in **Supplementary Fig. 8**. Overall, we found that a window of 200 SNPs and a jump of 1 were optimal for our datasets and enabled us to detect the well characterized selection candidates at the genome-wide significance threshold.

### Window size across epochs

Window sizes are also dependent on the number of segregating sites in a population as our windows are computed in units of SNPs. We chose to use a window size of 200 SNPs for all populations, after examining several population genetic parameters across different epochs (**Supplementary Fig. 8**). Importantly, the mean physical distance (bp) in a 200 SNP window *G12*_*ancient*_ window across epochs and number of segregating sites across epochs were quite consistent across epochs.

### Quality control for removing false sweeps

After running the selection scans and computing *G12*_*ancient*_ at each focal SNP, we performed quality control to remove spurious peaks that could have occurred due to artifacts or issues with the data. One reason a certain genomic position might have artificially high *G12*_*ancient*_ values is if the focal SNP and the SNPs within its window range overlap with regions of low recombination rate in the genome. The first step in post-processing/ quality control in our pipeline was to remove all windows with mean per-window recombination rates in the lowest fifth percentile genome-wide. Second, we also removed windows where the mean fraction of missing individuals (i.e., the mean of the fraction of missing individuals per SNP for all the SNPs in that window) was greater than the 70th percentile of the mean fraction of missing individuals for all windows. Third, our ascertainment scheme on the aDNA array results in each window having variable physical distance. While most windows are of similar length, some windows are in sites where the distance between positions is considerably lower or higher than the average. In order to show that our post-filtered data is largely unaffected by these issues, we regressed *G12*_*ancient*_ values against window size (measured in the physical distance), missingness, and recombination rate after the percentile-based removal process. We saw that the overall variability in the data explained by these three variables combined was less than 5%, suggesting that we had effectively removed their association with *G12*_*ancient*_ values (**Supplementary Table 4**). A final issue could be that there are windows where neighboring SNP positions are not captured well by the probes in our ascertainment scheme, and missingness rates are clustered even though the overall missingness rate is similar to other windows. To deal with these issues, we also removed windows that were consistently in the top 20 peaks genome-wide across a set of modern (the CEU, YRI, and STU populations) and the four ancient European populations we analyzed. The rationale for this is that it is quite unlikely that we see the same selective sweep across populations of such different ancestry, and across such a broad range of time and signals of that nature are highly likely to be due to data processing issues.

### Peak calling and gene annotation

As our main statistic is a multi-locus genotype-based scan, loci thought to be under selection lie in windows around top-scoring SNPs where the score (*G12*_*ancient*_ statistic value) is high compared to the rest of the genome. One issue with directly using the *G12*_*ancient*_ statistic value at each position to identify SNPs that appear to be selected significantly genome-wide is that many signals of selection at the SNP level are correlated due to LD. We wished to avoid identifying multiple high-scoring SNPs that are in linkage, as they might represent the same adaptive event. In order to account for this, we utilized a greedy clumping algorithm that looks for immediate positions upstream and downstream of a target SNP above a given threshold (https://github.com/ngarud/SelectionHapStats) as possible candidates. We assigned peaks to genes by taking the focal SNP in each peak and running Ensembl Variant Effect Predictor (VEP) ^68^ and annotated all protein-coding genes within 265kb distance upstream/ downstream of the target SNP and assigned the closest protein-coding gene for target SNP while annotating the *G12*_*ancient*_ peaks. The results of our analysis per epoch are shown in **Fig. 4a**.

On the 1.2 million positions captured on our array, we also annotated 47,384 as ‘potentially functional’ sites^8^ that lie in categories that overlap for certain SNPs. 1,290 SNPs were identified as targets of selection in Europeans by the Composite of Multiple Signals (CMS) test^69^; 21,723 SNPs identified as significant hits by genome-wide association studies, or with known phenotypic effect (GWAS); 1,289 SNPs with extremely differentiated frequencies between HapMap populations (HiDiff), 5,387 SNPs which tag HLA haplotypes and 13,672 expression quantitative trait loci (eQTLs). We then examined the distribution of *G12*_*ancient*_ statistic value across these categories of positions (**Fig. 4b**).

### Enrichment Analysis

We used the Functional Mapping and Annotation of Genome-Wide Association Studies tool to obtain significant gene sets for each epoch. The gene sets were produced by comparing the genes of interest against sets of genes from MsigDB using hypergeometric tests. We performed this analysis for gene sets from the GWAS and GO functional categories using FUMA^60^.

## Supporting information

Supplementary Data containing all analyzed samples

## Acknowledgments

We thank Arbel Harpak for useful discussions. Computing support for the project was supported on Director’s discretionary fund at the Texas Advanced Computing Cluster.

## Author contributions

D.P, M.H, N.R.G. and V. M. N. wrote the paper. D.P, M.H, N.R.G. and V.M.N. performed analysis.

## Data and materials availability

Code used for running *G12*_*ancient*_ selection scans can be found here: https://github.com/ngarud/SelectionHapStats, Code for running simulations can be found here: https://github.com/mariharris/Ancient_DNA_simulations. The ancient genomes used in this work can be accessed at Allen Ancient DNA Resource (AADR), version 51: https://reich.hms.harvard.edu/allen-ancient-dna-resource-aadr-downloadable-genotypes-present-day-and-ancient-dna-data.

---

**Supplementary Fig. 1:**
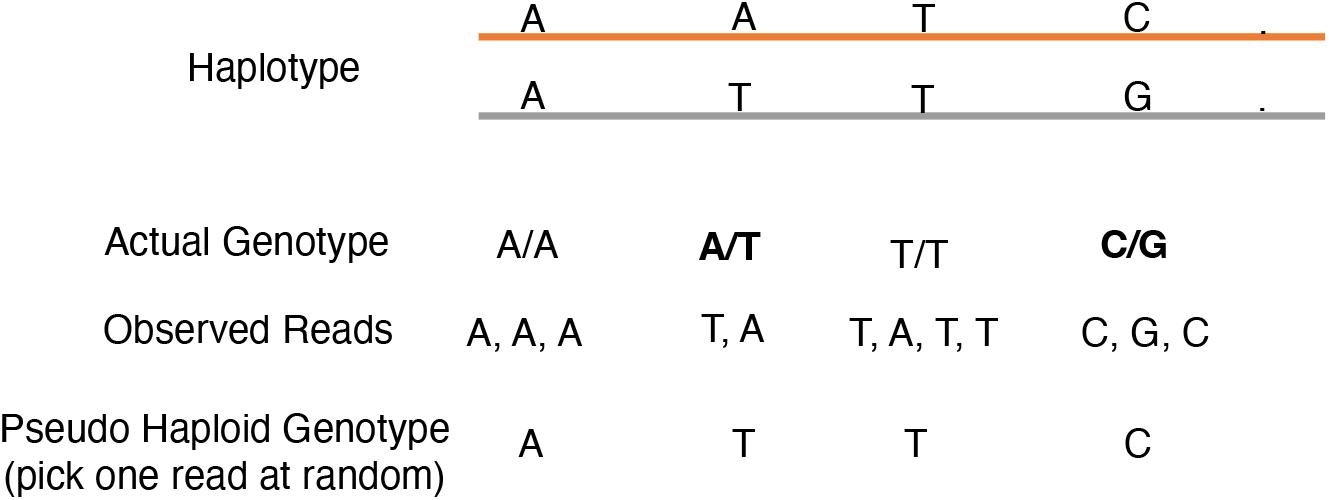
Pseudo haploidization scheme showing random allele calling for the generation of multi-locus genotypes.

**Supplementary Fig. 2:**
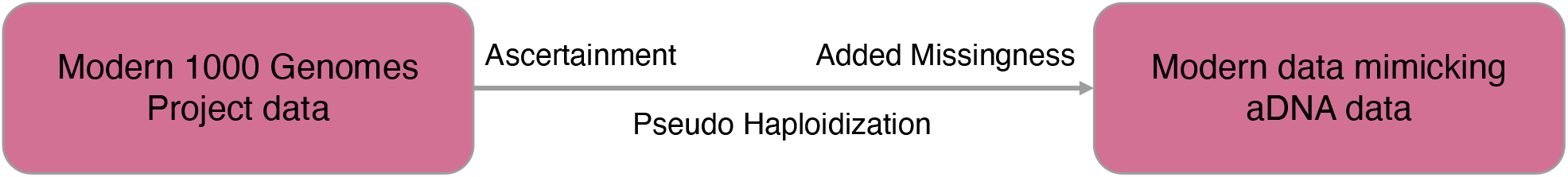
Data processing scheme, we take modern genomic data and apply ascertainment, pseudo-haploidization and add missingness to the data to make it mimic the artefacts of aDNA data used in this study.

**Supplementary Fig. 3:**
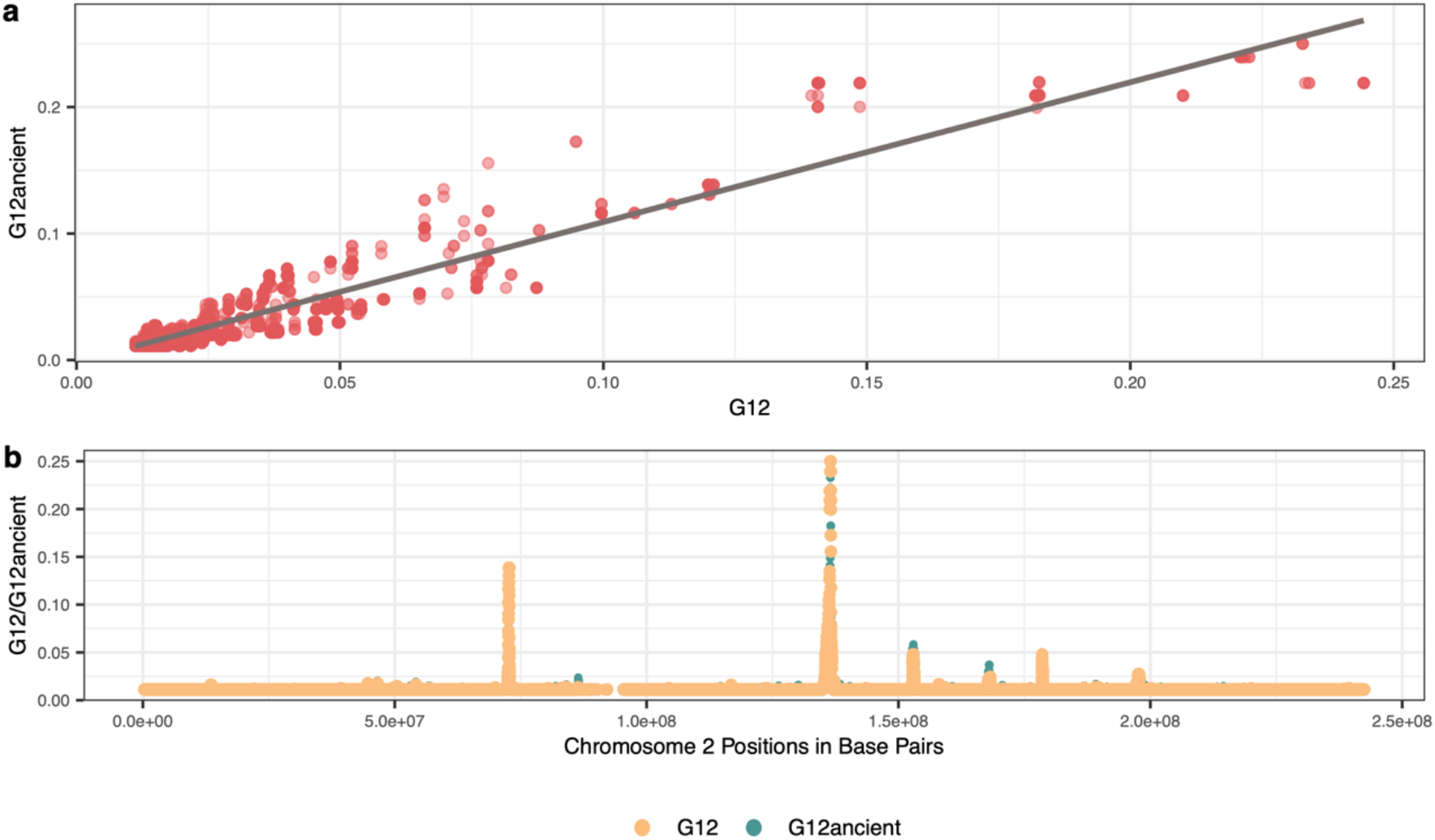
Plots showing strong positive correlation between *G12* and *G12*_*ancient*_ values for GBR individuals. **a** Scatter plot between *G12* and *G12*_*ancient*_ values with a line of best fit showing values are highly correlated. **b** Scatter plot between SNP positions and *G12* / *G12*_*ancien*t_ values showing that both plots overlay each other to a very higher degree.

**Supplementary Fig. 4:**
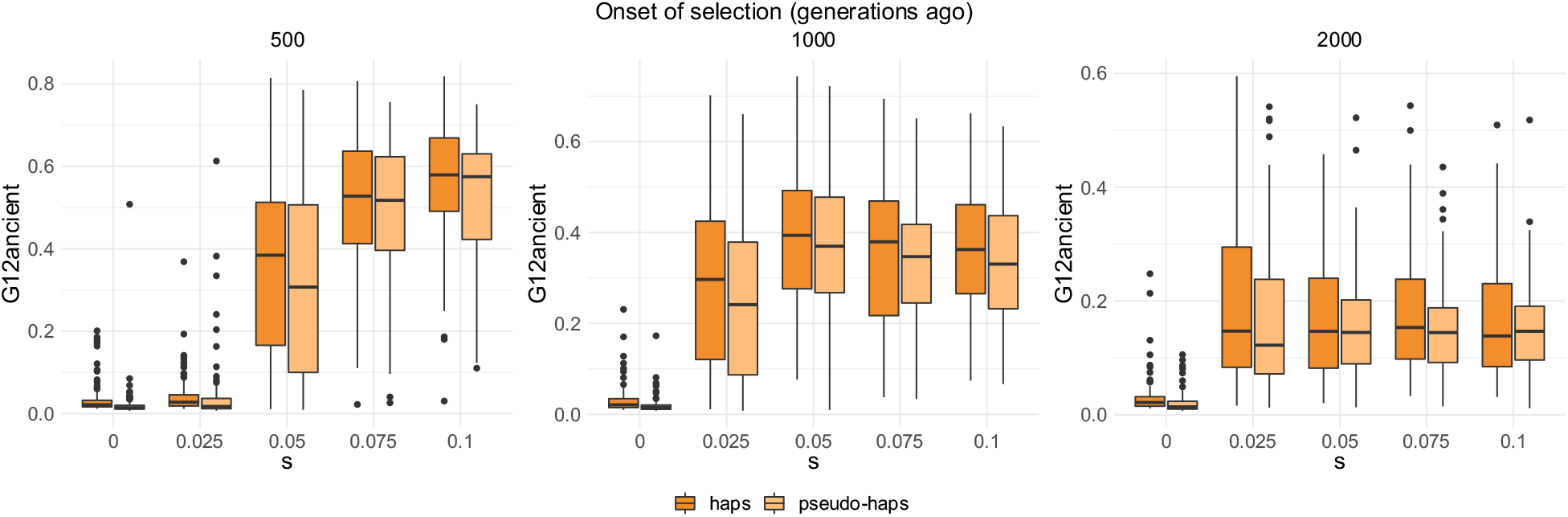
*G12* and *G12*_*ancient*_ values for 177 individuals sampled 40 generations ago. No missing data was added to the simulated data. We ran a total of 100 hard sweep simulations for each combination of parameters with mutation rate μ= 1.25×10^−8^ /bp, chromosome length L=5×10^5^ and recombination r = 5×10^−9^ events/bp.

**Supplementary Fig. 5:**
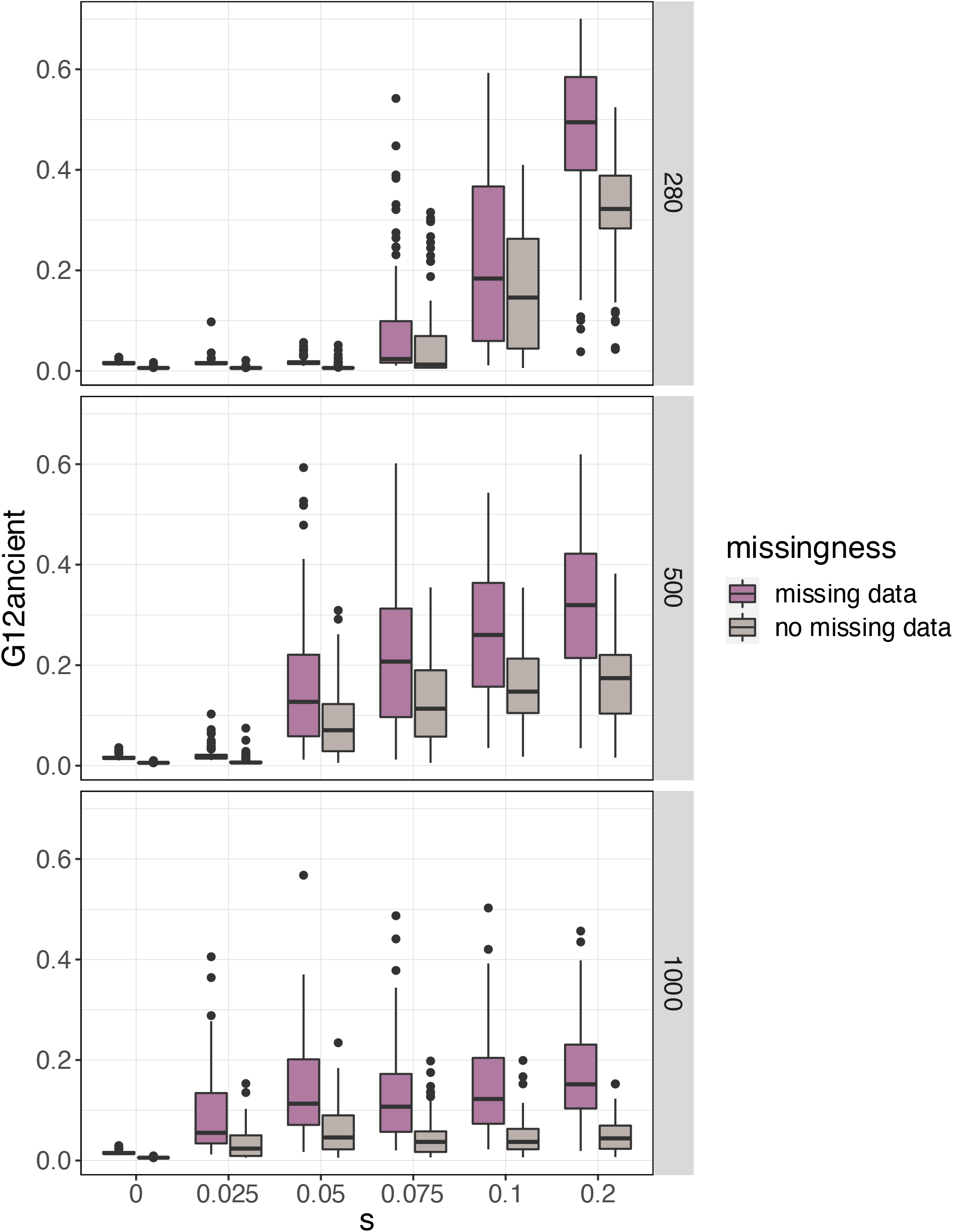
*G12*_*ancient*_ values for pseudo-haploidized simulated data from 177 individuals sampled 40 generations ago for a hard sweep model with mean rate of 0.55 missingness per SNP and a standard deviation of 0.23.

**Supplementary Fig. 6:**
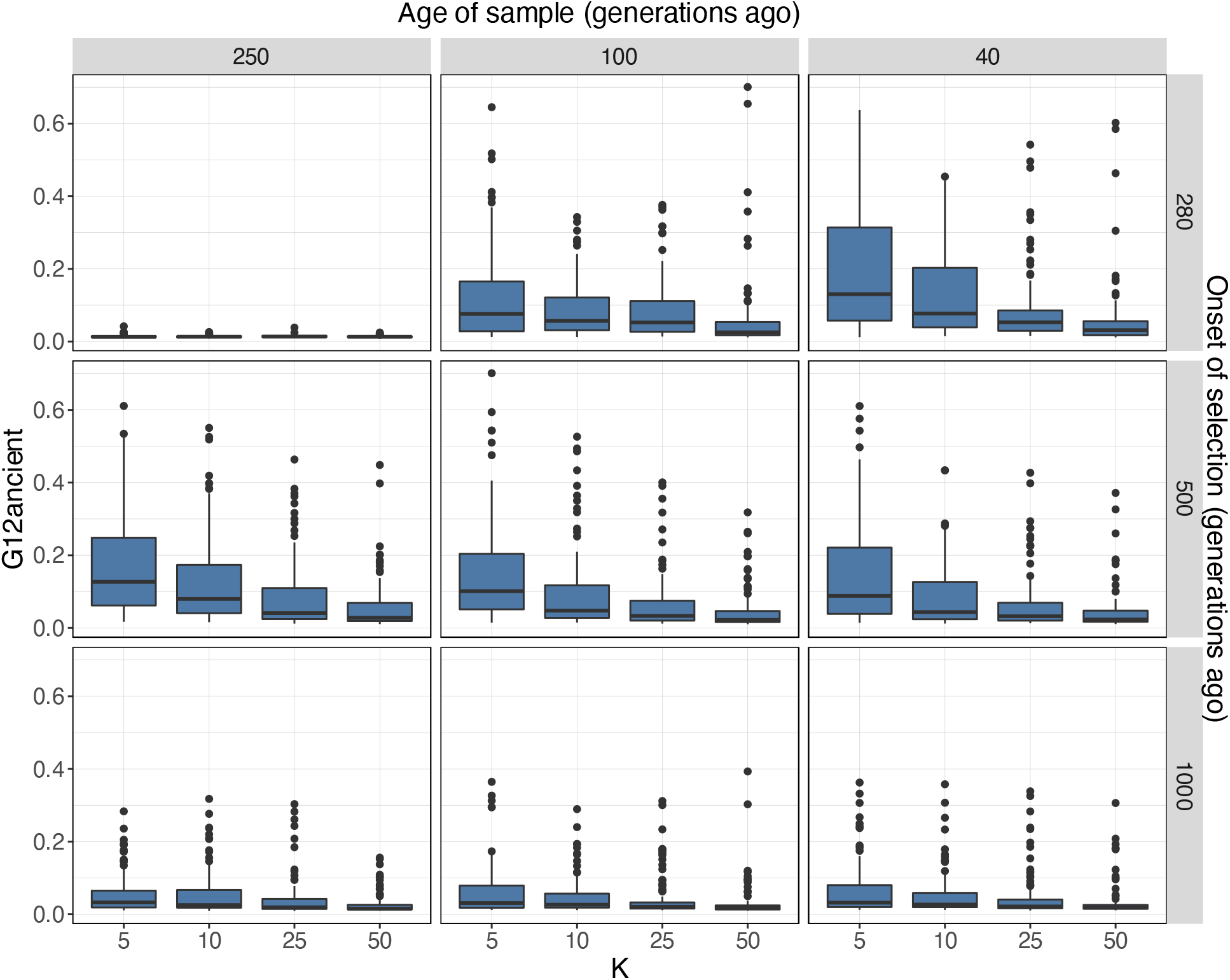
*G12*_*ancient*_ values in a soft sweep model. We introduced *K* beneficial mutations at the time of the onset of selection (rows), for *K=5,10,25* and *50*, where the higher *K* the softer the sweep. We sampled the population at 3 different time points (columns). We ran a total of 100 simulations for each combination of parameters with mutation rate μ = 1.25×10^−8^ /bp, chromosome length L = 5×10^5^, recombination r = 5×10^−9^ events/bp and s = 0.1. K = 0 corresponds to the scenario with no selection.

**Supplementary Fig. 7:**
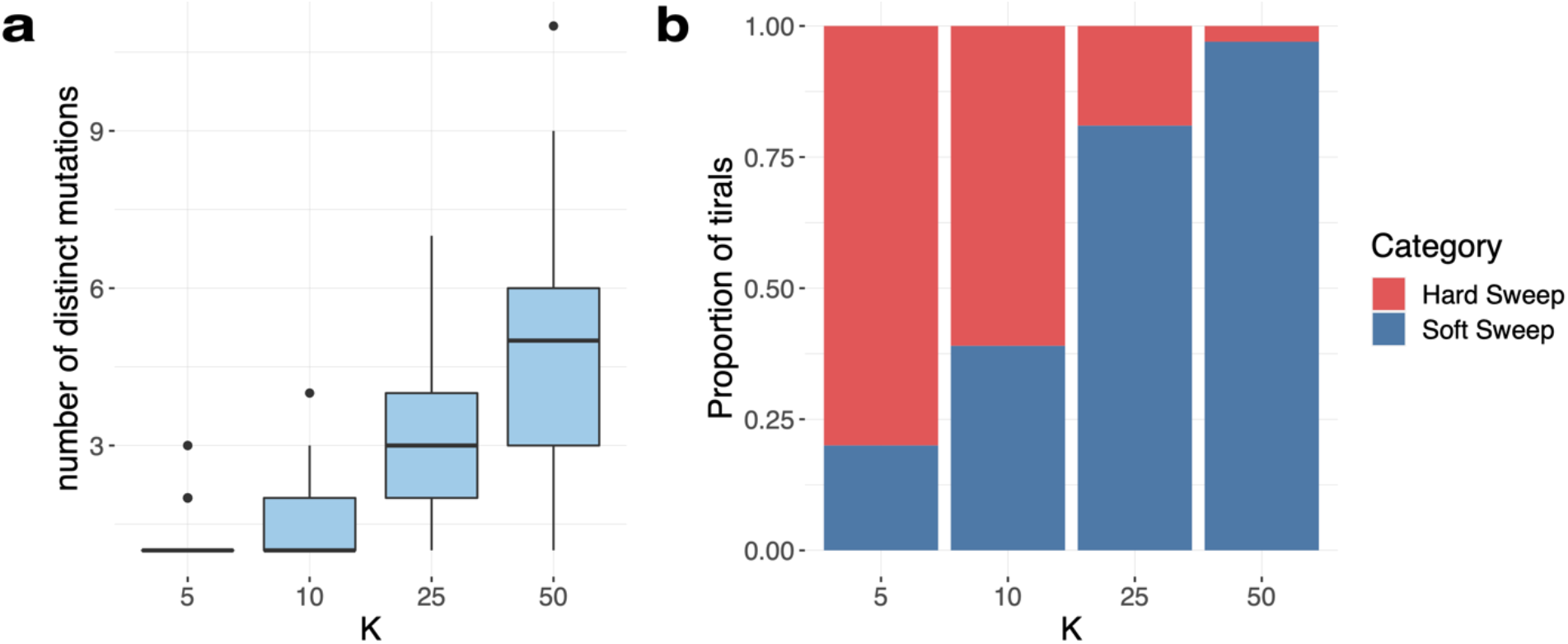
Softness of sweeps starting with *K* distinct mutations introduced 500 generations ago and sampled 40 generations ago. **a** Number of distinct mutations at the time of sampling. **b** Proportion of hard and soft sweeps as a function of *K*.

**Supplementary Fig. 8:**
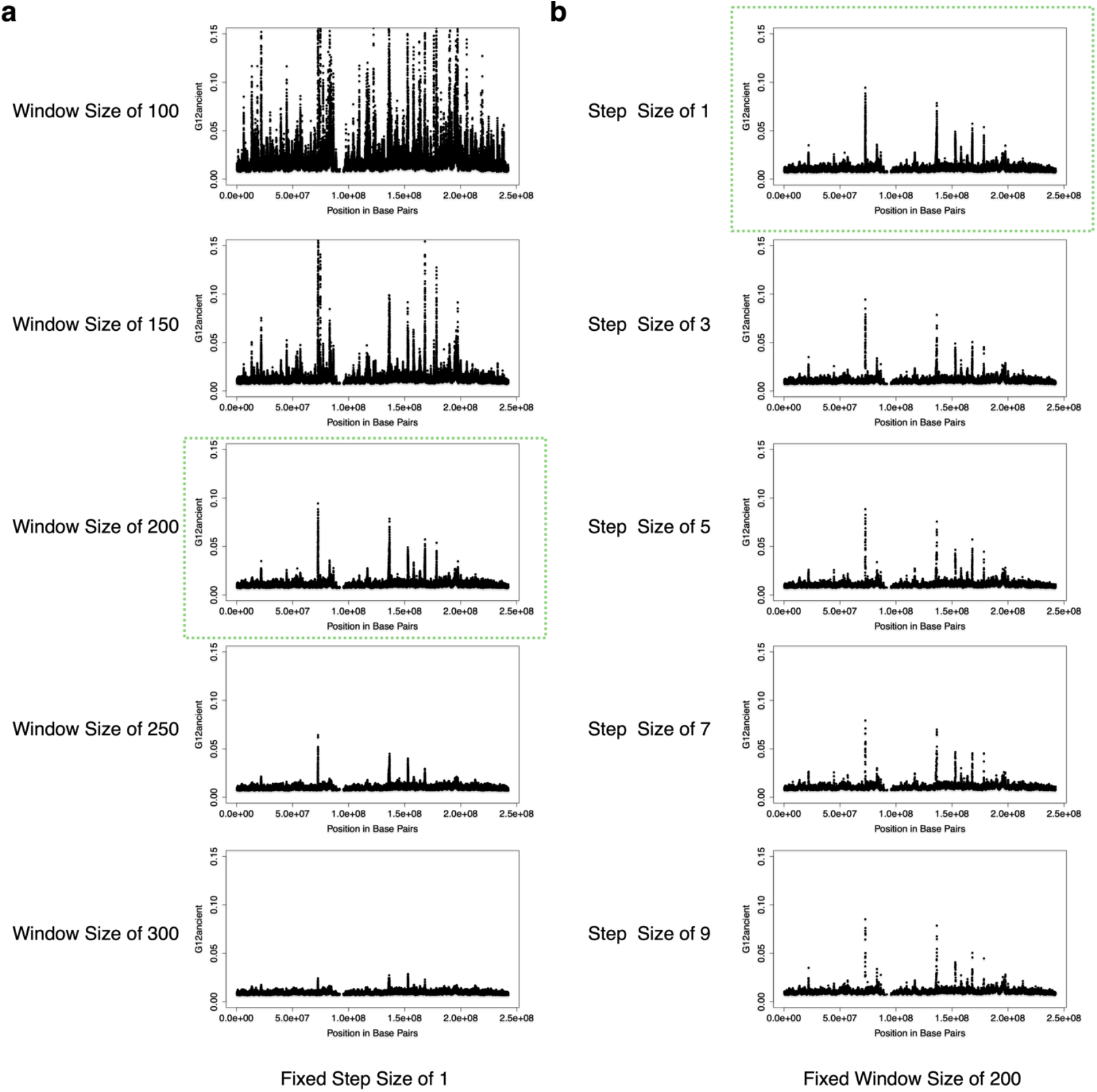
The performance of *G12*_*ancient*_ selection scans on different window size and step size values. **a** Variation of window size parameter while keeping step size fixed at 1, we observe window size of 200 as smaller window size resulted in inflated *G12*_*ancient*_ values and larger window size resulted in smaller *G12*_*ancient*_ values. **b** Variation of step size while keeping window size of 200, we observe that as we increase the step size, we lose a greater number of SNPs considered for calculation *G12*_*ancient*_ statistic and it results in loss of SNP density, so we fixed the step size as 1.

**Supplementary Table 1:** Differences between the mean fraction of missing individuals per SNP in modern samples vs. the ancient samples, pre, and post-data processing.

| Parameter | Modern Samples | Ancient Samples |
| --- | --- | --- |
| Mean Missingness (Preprocessing) | 0.0131 | 0.54827 |
| Mean Missingness (Post-processing) | 0.53529 | 0.54827 |

**Supplementary Table 2:**
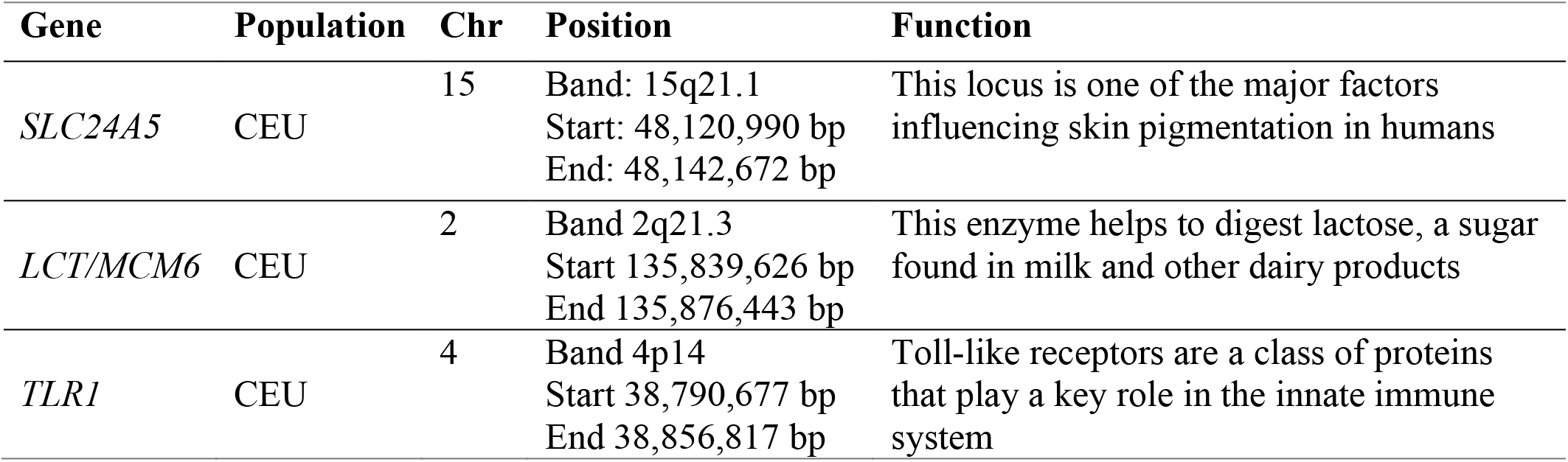
The variants of interest that are shown to be under selection by multiple natural selection studies on European genomes.

| Gene | Population | Chr | Position | Function |
| --- | --- | --- | --- | --- |
| *SLC24A5* | CEU | 15 | Band: 15q21.1<br>Start: 48,120,990 bp<br>End: 48,142,672 bp | This locus is one of the major factors influencing skin pigmentation in humans |
| *LCT/MCM6* | CEU | 2 | Band 2q21.3<br>Start 135,839,626 bp<br>End 135,876,443 bp | This enzyme helps to digest lactose, a sugar found in milk and other dairy products |
| *TLR1* | CEU | 4 | Band 4p14<br>Start 38,790,677 bp<br>End 38,856,817 bp | Toll-like receptors are a class of proteins that play a key role in the innate immune system |

**Supplementary Table 3:** A table showing the nucleotide diversity calculated for each epoch on a 200 SNP window. We used the vcftools --window-pi option which measures the nucleotide diversity in windows, with the number provided as the window size. We also show the number of segregating sites per base pair.

| Epoch | ND across 200<br>SNP window | Total<br>number of<br>sites | Segregating<br>sites | S/BP | Mean Window<br>Length (bp) |
| --- | --- | --- | --- | --- | --- |
| N | 0.00002752173 | 1233013 | 930906 | 0.754984 | 454774 (± 386740) |
| BA | 0.00002817481 | 1233013 | 953594 | 0.773385 | 454507 (± 386870) |
| IA | 0.00004058848 | 1233013 | 962723 | 0.780789 | 455005 (± 386849) |
| H | 0.00004152787 | 1233013 | 943001 | 0.764794 | 454605 (± 386845) |

**Supplementary Table 4:** Relationship between parameter choice and *G12*_*ancient*_ value suggests that overall *G12*_*ancient*_ statistics are unaffected by our choice of parameters.

| Variable | $G12_{ancient}$ | |
| --- | --- | --- |
|  | R<sup>2</sup> Score | Correlation Coefficient |
| Window Size | 0.013 | 0.1030 |
| Recombination Rate | 0.001 | -0.0261 |
| Missingness | 0.027 | 0.1634 |

## References

1. Vitti, J. J., Grossman, S. R. & Sabeti, P. C. Detecting natural selection in genomic data. Annu Rev Genet 47, 97–120 (2013).

2. Souilmi, Y. et al. Admixture has obscured signals of historical hard sweeps in humans. Nature Ecology & Evolution 2022 6:12 6, 2003–2015 (2022).

3. Ju, D. & Mathieson, I. The evolution of skin pigmentation-associated variation in West Eurasia. Proc Natl Acad Sci U S A 118, e2009227118 (2020).

4. Allentoft, M. E. et al. Population genomics of Bronze Age Eurasia. Nature 2015 522:7555 522, 167–172 (2015).

5. Mathieson, I. & Terhorst, J. Direct detection of natural selection in Bronze Age Britain. bioRxiv 2022.03.14.484330 (2022) doi:10.1101/2022.03.14.484330.

6. Gamba, C. et al. Genome flux and stasis in a five millennium transect of European prehistory. Nature Communications 2014 5:1 5, 1–9 (2014).

7. Lazaridis, I. et al. Ancient human genomes suggest three ancestral populations for present-day Europeans. Nature 2014 513:7518 513, 409–413 (2014).

8. Mathieson, I. et al. Genome-wide patterns of selection in 230 ancient Eurasians. Nature 2015 528:7583 528, 499–503 (2015).

9. Mathieson, I. et al. The genomic history of southeastern Europe. Nature 2018 555:7695 555, 197–203 (2018).

10. Sikora, M. et al. The population history of northeastern Siberia since the Pleistocene. Nature 2019 570:7760 570, 182–188 (2019).

11. Margaryan, A. et al. Population genomics of the Viking world. Nature 2020 585:7825 585, 390–396 (2020).

12. Mathieson, S. & Mathieson, I. FADS1 and the Timing of Human Adaptation to Agriculture. Mol Biol Evol 35, 2957–2970 (2018).

13. Le, M. K. et al. 1,000 ancient genomes uncover 10,000 years of natural selection in Europe. bioRxiv 2022.08.24.505188 (2022) doi:10.1101/2022.08.24.505188.

14. Sabeti, P. C. et al. Detecting recent positive selection in the human genome from haplotype structure. Nature 419, 832–837 (2002).

15. Voight, B. F., Kudaravalli, S., Wen, X. & Pritchard, J. K. A Map of Recent Positive Selection in the Human Genome. PLoS Biol 4, e72 (2006).

16. Garud, N. R., Messer, P. W., Buzbas, E. O. & Petrov, D. A. Recent Selective Sweeps in North American Drosophila melanogaster Show Signatures of Soft Sweeps. PLoS Genet 11, e1005004 (2015).

17. Garud, N. R. & Rosenberg, N. A. Enhancing the mathematical properties of new haplotype homozygosity statistics for the detection of selective sweeps. Theor Popul Biol 102, 94–101 (2015).

18. Ferrer-Admetlla, A., Liang, M., Korneliussen, T. & Nielsen, R. On Detecting Incomplete Soft or Hard Selective Sweeps Using Haplotype Structure. Mol Biol Evol 31, 1275–1291 (2014).

19. Martin, M. et al. WhatsHap: fast and accurate read-based phasing. bioRxiv 085050 (2016) doi:10.1101/085050.

20. Szpiech, Z. A. selscan 2.0: scanning for sweeps in unphased data. bioRxiv 2021.10.22.465497 (2022) doi:10.1101/2021.10.22.465497.

21. Kern, A. D. & Schrider, D. R. diploS/HIC: An Updated Approach to Classifying Selective Sweeps. G3 (Bethesda) 8, 1959–1970 (2018).

22. Harris, A. M., Garud, N. R. & Degiorgio, M. Detection and Classification of Hard and Soft Sweeps from Unphased Genotypes by Multilocus Genotype Identity. Genetics 210, 1429–1452 (2018).

23. Smith, A. J. Century-scale Holocene processes as a source of natural selection pressure in human evolution: Holocene climate and the Human Genome Project. http://dx.doi.org/10.1177/0959683607079003 17, 689–695 (2016).

24. Spyrou, M. A. et al. The source of the Black Death in fourteenth-century central Eurasia. Nature 2022 606:7915 606, 718–724 (2022).

25. Souilmi, Y. et al. An ancient viral epidemic involving host coronavirus interacting genes more than 20,000 years ago in East Asia. Curr Biol 31, 3504-3514.e9 (2021).

26. Brace, S. et al. Ancient genomes indicate population replacement in Early Neolithic Britain. Nature Ecology & Evolution 2019 3:5 3, 765–771 (2019).

27. Fernandes, D. M. et al. The spread of steppe and Iranian-related ancestry in the islands of the western Mediterranean. Nature Ecology & Evolution 2020 4:3 4, 334–345 (2020).

28. Harney, É. et al. A minimally destructive protocol for DNA extraction from ancient teeth. Genome Res 31, 472–483 (2021).

29. Lipson, M. et al. Parallel palaeogenomic transects reveal complex genetic history of early European farmers. Nature 2017 551:7680 551, 368–372 (2017).

30. Mathieson, I. et al. Genome-wide patterns of selection in 230 ancient Eurasians. Nature 2015 528:7583 528, 499–503 (2015).

31. Narasimhan, V. M. et al. The formation of human populations in South and Central Asia. Science (1979) 365, (2019).

32. Novak, M. et al. Genome-wide analysis of nearly all the victims of a 6200 year old massacre. PLoS One 16, e0247332 (2021).

33. Olalde, I. et al. The genomic history of the Iberian Peninsula over the past 8000 years. Science (1979) 363, 1230–1234 (2019).

34. Olalde, I. et al. The Beaker phenomenon and the genomic transformation of northwest Europe. Nature 2018 555:7695 555, 190–196 (2018).

35. Papac, L. et al. Dynamic changes in genomic and social structures in third millennium BCE central Europe. Sci Adv 7, 6941–6966 (2021).

36. Patterson, N. et al. Large-scale migration into Britain during the Middle to Late Bronze Age. Nature 2021 601:7894 601, 588–594 (2021).

37. O’Sullivan, N. et al. Ancient genome-wide analyses infer kinship structure in an Early Medieval Alemannic graveyard. Sci Adv 4, (2018).

38. Villalba-Mouco, V. et al. Survival of Late Pleistocene Hunter-Gatherer Ancestry in the Iberian Peninsula. (2019) doi:10.1016/j.cub.2019.02.006.

39. Fu, Q. et al. An early modern human from Romania with a recent Neanderthal ancestor. Nature 524, 216 (2015).

40. Pennings, P. S. & Hermisson, J. Soft sweeps II--molecular population genetics of adaptation from recurrent mutation or migration. Mol Biol Evol 23, 1076–1084 (2006).

41. Hermisson, J. & Pennings, P. S. Soft sweeps: molecular population genetics of adaptation from standing genetic variation. Genetics 169, 2335–2352 (2005).

42. Auton, A. et al. A global reference for human genetic variation. Nature 2015 526:7571 526, 68–74 (2015).

43. Akbari, A. et al. Identifying the Favored Mutation in a Positive Selective Sweep. Nat Methods 15, 279 (2018).

44. Haller, B. C. & Messer, P. W. SLiM 3: Forward Genetic Simulations Beyond the Wright– Fisher Model. Mol Biol Evol 36, 632–637 (2019).

45. Tennessen, J. A. et al. Evolution and functional impact of rare coding variation from deep sequencing of human exomes. Science 337, 64–69 (2012).

46. Beleza, S. et al. The Timing of Pigmentation Lightening in Europeans. Mol Biol Evol 30, 24 (2013).

47. Gerbault, P. et al. Evolution of lactase persistence: an example of human niche construction. Philosophical Transactions of the Royal Society B: Biological Sciences 366, 863 (2011).

48. Segurel, L. et al. Why and when was lactase persistence selected for? Insights from Central Asian herders and ancient DNA. PLoS Biol 18, e3000742 (2020).

49. Evershed, R. P. et al. Dairying, diseases and the evolution of lactase persistence in Europe. Nature 2022 608:7922 608, 336–345 (2022).

50. Mathieson, I. & Terhorst, J. Direct detection of natural selection in Bronze Age Britain. bioRxiv 2022.03.14.484330 (2022) doi:10.1101/2022.03.14.484330.

51. Ju, D. & Mathieson, I. The evolution of skin pigmentation-associated variation in West Eurasia. Proc Natl Acad Sci U S A 118, e2009227118 (2020).

52. Donnelly, M. P. et al. A global view of the OCA2-HERC2 region and pigmentation. Hum Genet 131, 683 (2012).

53. Wilde, S. et al. Direct evidence for positive selection of skin, hair, and eye pigmentation in Europeans during the last 5,000 y. Proc Natl Acad Sci U S A 111, 4832–4837 (2014).

54. Chen, Y. et al. Variations in DNA elucidate molecular networks that cause disease. Nature 452, 429–435 (2008).

55. Becker, P. H. et al. Adenosine kinase deficiency: Three new cases and diagnostic value of hypermethioninemia. Mol Genet Metab 132, 38–43 (2021).

56. Li, H. et al. Hepatocyte Adenosine Kinase Promotes Excessive Fat Deposition and Liver Inflammation. Gastroenterology 164, 134–146 (2023).

57. Moser, E. K. & Oliver, P. M. Regulation of autoimmune disease by the E3 ubiquitin ligase Itch. Cell Immunol 340, (2019).

58. Yin, Q., Wyatt, C. J., Han, T., Smalley, K. S. M. & Wan, L. ITCH as a potential therapeutic target in human cancers. Semin Cancer Biol 67, 117–130 (2020).

59. Miyagawa, H. et al. Association of polymorphisms in complement component C3 gene with susceptibility to systemic lupus erythematosus. Rheumatology 47, 158–164 (2008).

60. Watanabe, K., Taskesen, E., van Bochoven, A. & Posthuma, D. Functional mapping and annotation of genetic associations with FUMA. Nature Communications 2017 8:1 8, 1–11 (2017).

61. Childebayeva, A. et al. Population Genetics and Signatures of Selection in Early Neolithic European Farmers. Mol Biol Evol 39, (2022).

62. Mallick, S. et al. The Allen Ancient DNA Resource (AADR): A curated compendium of ancient human genomes. (2023) doi:10.1101/2023.04.06.535797.

63. Fu, Q. et al. A revised timescale for human evolution based on ancient mitochondrial genomes. Curr Biol 23, 553–559 (2013).

64. Korneliussen, T. S., Albrechtsen, A. & Nielsen, R. ANGSD: Analysis of Next Generation Sequencing Data. BMC Bioinformatics 15, 1–13 (2014).

65. Price, A. L. et al. Principal components analysis corrects for stratification in genome-wide association studies. Nature Genetics 2006 38:8 38, 904–909 (2006).

66. Patterson, N., Price, A. L. & Reich, D. Population Structure and Eigenanalysis. PLoS Genet 2, e190 (2006).

67. Furtwängler, A. et al. Ancient genomes reveal social and genetic structure of Late Neolithic Switzerland. Nature Communications 2020 11:1 11, 1–11 (2020).

68. McLaren, W. et al. The Ensembl Variant Effect Predictor. Genome Biol 17, 1–14 (2016).

69. Grossman, S. R. et al. Identifying Recent Adaptations in Large-scale Genomic Data. Cell 152, 703 (2013).

